# Design and Immunogenicity of SARS-CoV-2 DNA vaccine encoding RBD-PVXCP fusion protein

**DOI:** 10.1101/2022.10.06.511104

**Authors:** Dmitri Dormeshkin, Mikalai Katsin, Maria Stegantseva, Sergey Golenchenko, Michail Shapira, Simon Dubovik, Dzmitry Lutskovich, Anton Kavaleuski, Alexander Meleshko

## Abstract

The potential of immune evasive mutations accumulation of the SARS-CoV-2 virus has led to its rapid spread causing over 600 million confirmed cases and more than 6.5 million confirmed deaths. Huge demand for the rapid development and deployment of low-cost and effective vaccines against emerging variants renews interest in DNA vaccine technology. Here we report a rapid generation and immunological evaluation of novel DNA vaccine candidates against Wuhan-Hu-1 and Omicron variants, based on the RBD protein fused with the Potato virus X coat protein (PVXCP). Delivery of DNA vaccines using electroporation in a two-doses regimen induced high antibody titers and profound cellular response in mice. Antibody titers induced against Omicron variant of the vaccine were sufficient for the effective protection against both the Omicron and Wuhan-Hu-1 virus infections. PVXCP protein in the vaccine construct shifted immune response to the favorable Th1-like type and provided oligomerization of RBD-PVXCP protein. A naked DNA delivery by the needle-free injection device allowed us to achieve antibody titers comparable with the mRNA-LNP delivery in rabbits. This data identifies the RBD-PVXCP DNA vaccine platform as a promising solution for robust and effective SARS-CoV-2 protection, supporting further translational study.

## 1 Introduction

The current ongoing global pandemic caused by severe acute respiratory syndrome coronavirus 2 (SARS-CoV-2) has resulted in more than 6.5 million coronavirus disease (COVID-19)-related deaths as of September 2022 (www.worldometers.info/coronavirus/).

SARS-CoV-2 is a new pathogenic human coronavirus, which also includes hCoV-OC43, hCoV-HKU1, hCoV-229E, hCoVNL63 and two life threatening beta-coronaviruses: Middle East respiratory syndrome coronavirus (MERS-CoV) and SARS-CoV^1^. The viral genome is represented by a positive-sense, single-stranded RNA encoding four major structural proteins: spike (S) in a form of homotrimer, nucleocapsid (N), membrane (M) and envelope (E)^2^. S protein is cleaved at a polybasic cleavage site at its S1/S2 junction into S1 and S2 subunits during their biosynthesis in the infected cells by proprotein convertases such as furin. S protein of the mature virion consists of these two non-covalently associated subunits^3^.

The entry of SARS-CoV-2 begins with the binding of receptor binding protein (RBD) within the S1 subunit of S protein to angiotensin-converting enzyme 2 (ACE2) on a host cell, leading to conformational changes, S1 shedding and an additional S2 cleavage site exposition. Later, transmembrane protease serine 2 (TMPRSS2) at the cell surface or cathepsin L in the endosomal compartment operates cleavage at S2 site, releasing the fusion peptide and subsequent initiating of a fusion pore formation^4^. The position switching of the RBD from down to up executes receptor binding while up to down assists the virus to impede the immune surveillance. The availability of the receptor-binding determining region to the ACE2 is controlled by the hinge-like conformational motion of the RBD^5^. A mutation at the 614 amino acid i.e. Asp614-Gly, being present in all mutant strains and absent in the wild type Wuhan-Hu-1 strain, has been reported to enhance the up conformation of the RBD and diverse epitopes accessibility. Such configuration makes the virus more infectious and susceptible to neutralizing antibodies^6^. The last fact could be taken into account in the S-protein based SARS CoV-2 vaccine design development, because closed S proteins could expose other conformational epitopes eliciting different antibody specificity responses^7^.

N and M proteins, as well as Nsp8 and ORF9b of SARSCoV-2 are engaged in escape from detection by innate immune sensors. SARS-CoV-2 N protein suppresses the activity of STAT1 and STAT2, interfering with the IFN signaling pathway^8^. Moreover, M protein is able to bind RIG-I, MDA5, MAVS, and TBK1 to prevent their interaction suppressing type I and III IFN production^9^. Nsp8 and ORF9b proteins also suppress the IFN signaling pathway by direct binding to MDA5 CARD and translocator of outer membrane 70, respectively^10,11^. These defects in innate immunity caused by SARS CoV-2 could impair humoral and cellular immunity acquisition. In one study, it has been shown that serum from vaccinated patients had 17 times higher neutralization capacity than serum collected from patients after natural infection^12^.

Effective vaccines have been developed and globally introduced, including inactivated whole virus vaccines, protein/peptide subunit vaccines, nanoparticle vaccines, vector vaccines, and nucleic-based formulations (mRNA and DNA vaccines), each having its own pros and cons^13,14,15^.

Despite the unprecedented success in SARS-CoV-2 vaccine development, incorporating S protein from wild type Wuhan-Hu-1 strain, novel strains of SARS-CoV-2 have emerged facilitating mild to moderate escape from humoral immunity, which includes but not limited to B.1.1.7 (UK; alpha variant), B.1.351 (SA; beta variant), P.1 (Brazil; gamma variant), B.1.617.2 (India; delta variant) and recently Omicron (B.1.1.529) with its derivatives. Several studies reported a 2-to 4-fold reduction in neutralization against Delta, a 6-fold or higher reduction in neutralization against Beta, and a 29-fold reduction in neutralization against Omicron for both convalescent and vaccinated individuals^16,17,18^. Despite reduced vaccine efficacy against mutant strains, effectiveness remained high against hospitalization or severe disease^19^.

Omicron has 32 mutations in the S-protein and 15 mutations are located in the RBD, which is the main key for viral-cell interactions and entry mediated by ACE-2^20,21^. Serum from vaccinated patients or convalescents from Alpha (B.1.1.7), Beta (B.1.351), Delta (B.1.617.2) SARS-CoV-2 infection has substantially decreased neutralization capacity of Omicron subvariant as well as therapeutic antibodies^22,23^. An additional prolonged antigen stimulation by vaccine boost or infection can increase the neutralization breadth and resistance to RBD mutations, for instance against Omicron, that is reflected by an ongoing germinal center reaction in RBD-specific memory B-cells with somatic hypermutations acquisition, affinity maturation, and at the same time contraction of humoral responses to other SARS-CoV-2 domains^24,25,26^. It has been shown, that homotypic nanoparticle RBD SARS CoV-2 vaccine induced broad cross-reactive neutralization and binding to zoonotic coronaviruses (SHC014, WIV1, Yun 11, BM-4831 and BtKY72), in which RBD sequence homology varied from 68% to 95%. Such results were not reproduced by soluble S-trimeric vaccine and convalescent serum^27^.

RBD, NTD and S2 are the main domains of SARS-CoV2 which could induce neutralizing antibodies^28,29^. SARS-CoV-2 RBD is the main immunodominant site of S protein for neutralizing antibodies induction. It has been shown that depletion of anti-RBD antibodies in convalescent patient sera resulted in the loss of more than 90% of neutralizing activity of the sera against SARS-CoV-2^30^.

RBD is considered a low immunogenic antigen due to the relatively small length of antigen and strategies to promote immunogenicity, including the use of appropriate adjuvants or the addition of exogenous sequences capable of potentiating immune responses should be discussed in vaccine development ^31^. Previous studies revealed that vaccines incorporating SARS-CoV full-length S protein could induce harmful immune responses with enhanced infection or liver damage after virus challenge that is not seen with RBD SARS-CoV vaccines^32,33,34^. Implicating RBD antigen for SARS CoV-2 vaccine development instead of full S-protein could potentially decrease the probability of antibody-dependent enhancement (ADE) development as well as induce stronger humoral and cellular immune responses^35,36,37^.

Despite the fact that mRNA vaccines became one of the breakthroughs in COVID-19 vaccine development, DNA vaccines are gaining more and more attention. DNA vaccines have not reached their full potential yet and have a wide window of improvement, including DNA delivery, *in vivo* expression enhancement and vaccine design improvement. There are several advantages of DNA vaccines: an unlimited number of reapplications due to lack of immune response against the vector, natural Th1 response skewing, flexibility in vaccine upgrade, simplicity and rapidity to make and scale up production with a resultant affordable price for the product. Furthermore, DNA vaccines proved to be safe and not reactogenic, possess high stability and have no need for the sophisticated low-temperature supply chain^38^.

The MERS DNA vaccine has been tested in a Phase I trial, where seroconversion occurred in 61 (94%) of 65 participants after two and three vaccinations. T-cell responses were detected in more than 71% percent of patients. Response tended to be stable: at week 60, vaccine-induced humoral and cellular responses were detected in 51 (77%) of 66 participants and 42 (64%) of 66, respectively^39^.

Most typically, DNA vaccines yield low immune responses, mainly due to the inefficiency of DNA delivery into macroorganisms. Improved delivery by electroporation and needle-free injectors (for example PharmaJet Needle-free Injection System) increased in vivo antigen expression and immune responses to DNA vaccines. Other approaches such as reduction of the size of the DNA plasmid, deletion of antibiotic resistance genes, and addition of immunostimulatory sequences into the backbone of the DNA vaccine have been shown to increase the immunogenicity of SARS-COV-2 DNA vaccine^40^.

But there is an additional window of DNA vaccine improvement^41, 42, 43^. To deepen the humoral immune response to our DNA SARS-COV-2 vaccine, we implemented several solutions: I) fusion of SARS-CoV-2 antigen to Potato virus X coat protein (PVXCP) to ensure an assembly of VLP and Th1 immune response skewing: II) selection RBD as an antigen of SARS-COV-2 to enhance the protection from VOC’s and ensure the safety of our SARS-CoV-2 vaccine.

Based on aforementioned arguments, we developed different designs of SARS-CoV-2 DNA-vaccines incorporating RBD as a backbone antigen and compared their efficacy *in vivo*. The plasmid DNA vaccine candidate, encoding Omicron BA.1 SARS-CoV-2 RBD and PVXCP demonstrated the high antibody response and strong Th1-biased cellular response. It was demonstrated that the delivery of the DNA vaccine RBD-PVXCP by needle-free injection system induced >300 000 RBD-specific IgG mean endpoint titer in rabbits. Our results thus identify RBD-PVXCP as a robust and effective vaccine candidate against SARS-CoV-2.

## 2 Material and Methods

### 2.1 Molecular Dynamics simulation

The spatial structures of RBD and HR2 domain were obtained from PDB (PDB IDs: 7KLW and 6XR8 respectively), as well as a peculiarities of RBD-HR2 domain joint^44,45^. For the construction of the 3-dimension structure of PVXCP PDB structure with PDB ID 6R7G was used^46^. Three protein domains were connected to each other into one chimeric protein structure taking into the account allowed values for φ and ψ angles (various resulting structures were additionally checked using Molecular Modelling Toolkit python library).

For further structure optimization of chimeric protein molecular dynamics (MD) the minimization, heating, equilibration, and free dynamics stages were performed. The minimization protocol included 4500 steps using the gradient descent method and 500 steps using the conjugate gradient method. Heating was carried out for 1 ns to a temperature of 298.15 K (NVT ensemble). At the next step, the simulated system was equilibrated for 1 ns at 298.15 K at constant pressure. Simulation of free dynamics was carried out on a time interval of 500 ns (NPT ensemble). A constant pressure in the system was maintained using an external barostat (relaxation time, 2 ps). A constant temperature was maintained using a Langevin thermostat (collision frequency, 2 ps^−1^). At all stages of modeling, the cutoff value was taken equal to 8.0 Å. The calculation was carried out in an explicit solvent (water, TIP3P model, the size of the modeling area was 8.0 Å from the protein surface).

After MD, optimized structure of chimeric molecule RBD-HR2-PVXCP was used for replacing the original monomers of the structure from 6R7G (one helix turn). Obtained multimolecular complex were analyzed by the previously described MD (50 ns simulation) in order to estimate the probability of complex dissociation.

### 2.2 Preparation of DNA constructs

Codon optimized synthetic sequences were generated by gene synthesis (Synbio Technologies, USA) and subcloned into mammalian expression vector pIF (Figure S1.). This vector was designed for DNA *in vivo* delivery and has a reduced size of a backbone (3.1 kb) with a minimal set of bacteria originated elements, including: pBR322 origin of replication, kanamycin resistance gene, CMV enhancer/promoter without intron A and bGH poly(A) signal. Transfection-grade plasmid DNA was isolated from XL10Gold cell overnight culture with the NucleoBond™ Xtra Midi Kit (Macherey-Nagel, Germany).

### 2.3 In Vitro DNA vaccines expression

All the plasmids for analytical expression were purified using PureLink™ HiPure Plasmid Miniprep Kit (#K210003, Invitrogen). Plasmid vectors were transiently transfected into HEK293 FreeStyle cells using polyethylenimine (PEI) transfection. 24 hours before transfection HEK 293 FreeStyle cell line was subcultured to final density 5*10^5^ cells per ml of antibiotic-free FreeStyle medium to reach 1*10^6^ cells per ml to the transfection day in a total volume of 2 ml. A DNA and PEI have been taken in amounts of 2 μg and 3.2 μg respectively per well. DNA:PEI complexes were made by mixing of separately prepared solutions of 2 μg of DNA and 3.2 μg of PEI diluted by OptiMem to final volumes of 37 μl. After PEI solution adding to DNA, the mixture rested at room temperature for 25 minutes. The final mixture was added to the cells followed by placing them in an incubator (37°C, 5% CO_2_) for 5 days with a constant shaking of 120 rpm. Conditioned expressing medium was centrifuged to remove cell debris and then cell medium was analyzed by Western Blotting using anti-S-protein polyclonal antibodies (#E-AB-V1006, Elabscience).

### 2.4 Identification of SARS-CoV-2 binders

Tris-glycine Novex gels (#EC6021BOX, ThermoFisher Scientific) were loaded with 10 *μ*l of cell supernatant on the 5th day after the transfection. Gels were run at 120 V for 1,5 hours in Tris-tricine-SDS buffer. After electrotransfer in the Towbin buffer using Trans-Blot Turbo Transfer System (Bio-Rad), PVDF membrane was blocked with 5% filtered skimmed milk in PBS overnight at 4°C. The membrane was washed and then incubated with 1:5 000 SARS-COV-2 Spike RBD Polyclonal Antibody antibodies (#E-AB-V1006, Elabscience) for 1 h at RT and with 1:10 000 Goat Anti-Rabbit-HRP (#31460, Invitrogen), for 1 h at RT. After washing, the membrane was developed using ECL Clarity Substrate (Bio-Rad, USA) and imaged using Azure C300 imager (Azure Biosystems, China).

### 2.5 Mice immunization

BALB/c female mice (6 weeks old) were purchased from the Rappolovo Animal Farm of the Russian Academy of Medical Sciences and housed in the animal facility of Institute of Physiology of the National Academy of Sciences of Belarus (Minsk, Belarus). For the vaccination purposes mice received 50 μL intramuscular (IM) injection of 2.5 μg or 50 μg of pDNA with the following electroporation (EP) in the tibialis muscle of the shaved hind leg on days 0 and 14 of the experiment. EP was performed with AgilePulse In Vivo System (BTX). Prior to the immunization all mice were anesthetized by isoflurane inhalation with RAS-4 Rodent Anesthesia System (PerkinElmer). Mice were euthanized on day 28 for terminal blood collection and spleens were harvested for cellular assays. All animal experiments were done according to the Belarusian Guide for the Care and Use of Laboratory Animals.

### 2.6 Rabbits immunization

Ten female New Zealand white rabbits were housed in the animal facility of Institute of bioorganic chemistry of the National Academy of Sciences of Belarus (Minsk, Belarus). Rabbits were divided into four groups:

- Group 1 (n=2). 100 μg pDNA (vector control - pIF) solution in PBS was administered intramuscular by using needle injection
- Group 2 (n=2). 100 μg pDNA (v1) solution in PBS was administered by intradermal needle injection followed by electroporation with AgilePulse In Vivo System (BTX).
- Group 3 (n=3) 100 μg pDNA (v1) solution in PBS was administered intramuscular by using needle injection.
- Group 4 (n=3) 100 μg pDNA (v1) solution in PBS was administered to the skin by using the needle-free injection system.

All rabbits were vaccinated on days 0 and 14 of the experiment. Rabbits from the Group 3 and Group 4 received an additional third boost of the vaccine on the 42nd day of the experiment. On days 28, 56, 75 and 105 blood was collected by the ear vein sampling.

### 2.7 ELISA

For endpoint titer determination a 96-well microtiter plate was coated with 2 μg/ml RBD in 100 μl PBS overnight at 4 °C. Afterward, the plate was blocked with 5% skimmed milk in PBS for 2 hours at RT. Serial dilutions of mice\rabbit serum were subjected to the wells in 2.5% skimmed milk in PBS and incubated at RT for 60 min. Animal immune serum samples were heated at 55°C for 30 minutes before use. After incubation, the wells were washed six times with 0.05% PBST and 100 μL of anti-IgG-HRP antibody conjugated (1:5 000 dilution) was added. After 1 h of incubation and five washes, positive binders were determined upon 100 μl TMB substrate addition. The absorbance at 450 nm was determined using Clariostar (BMG, Germany) reader after stopping the reaction by adding 100 μL of 2M H_2_SO_4_ per well.

Endpoint titers were calculated using in-house build Python script (https://github.com/MShapira?tab=overview&from=2022-09-01&to=2022-09-28). Algorithm allows to calculate cutoff value for each sample group and endpoint titer based on the estimated cutoff^47^. For the data approximation the further equation is used:

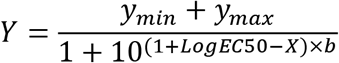

where:

*LogEC50* - is the serum dilution that gives a response half way between *y_min_* and *y_max_*,
*b* - describes the steepness of the curve.

This description of four-parameter logistic curve allows to approximate almost all experimental data except sets without at least one expressed plato (bottom or top).

### 2.8 ELISPot

Spleens from immunized mice were collected in sterile tubes containing RPMI-1640 (ThermoFisher Scientific) media supplemented with 10% fetal bovine serum and 2X Antibiotic-Antimycotic (ThermoFisher Scientific). The cell suspension was obtained by rubbing the spleen through a 100 μm cell strainer (Corning). The mononuclear fraction of cells was separated from erythrocytes by gradient centrifugation on Histopaque-1077 (Sigma-Aldrich).

Interferon gamma production was measured using the ELISpot Plus: Mouse IFN-γ (ALP) (Mabtech). Splenocytes were counted and 250,000 cells were plated per well into pre-covered plates and stimulated overnight with 10 μg/mL SARS-CoV-2 (S1) peptide pool (Mabtech) or medium as negative control or PHA 2 μg/mL as positive experimental control. The next day, plate was washed off, and treated with a biotinylated anti-IFN-γ detection antibody followed by a streptavidin-ALP conjugate resulting in visible spots. The plates were dried and the spots were counted on AID iSpot Spectrum (AID GmbH) and analyzed with EliSpot Software Version 7.x.

### 2.9 Statistical analysis

The statistical analysis was performed using GraphPad Prism software 8.4 (GraphPad Software, Inc. LA Jolla, CA, USA).

## 3 Results

### 3.1 SARS-CoV-2 DNA vaccine candidates design and in vitro analysis

A number of DNA-encoded vaccines were recently developed, but all of them have a common drawback of insufficient level of humoral responses in large animals^40,48,49^.

We hypothesized that oligomerization of DNA-encoded SARS-CoV-2 vaccine could enhance humoral response resulting in higher titers even using the same nucleic acid delivery technology and commonly used vectors. It also has been found that 97.9% recovered COVID-19 patients exerted high titer IgG specific for HR region, but only the IgG titer to RBD was able to differentiate recurrent viral RNA-positive from persistently RNA negative patients^50^.

We designed a number of DNA-constructs (Figure 1A) encoding SARS-CoV-2 RBD proteins: 1) monomeric RBD (v0), 2) RBD fused with the PVXCP fragment via rigid linker (RBD-PVXCP (v1)) and its variation with the N-terminal hexahistidine tag (His6-RBD-PVXCP(His6-v1)), 3) RBD-PVXCP protein with the HR2 region incorporated between them (RBD-HR2-PVXCP(v2)) and its variations with the N-terminal hexahistidine tag (His6-RBD-HR2-PVXCP(His6-v2)), 4) RBD-PVXCP with the RBD fragment harboring Omicron B.1.1.529 mutations (oRBD-PVXCP (v1.om)).

**Figure 1.**
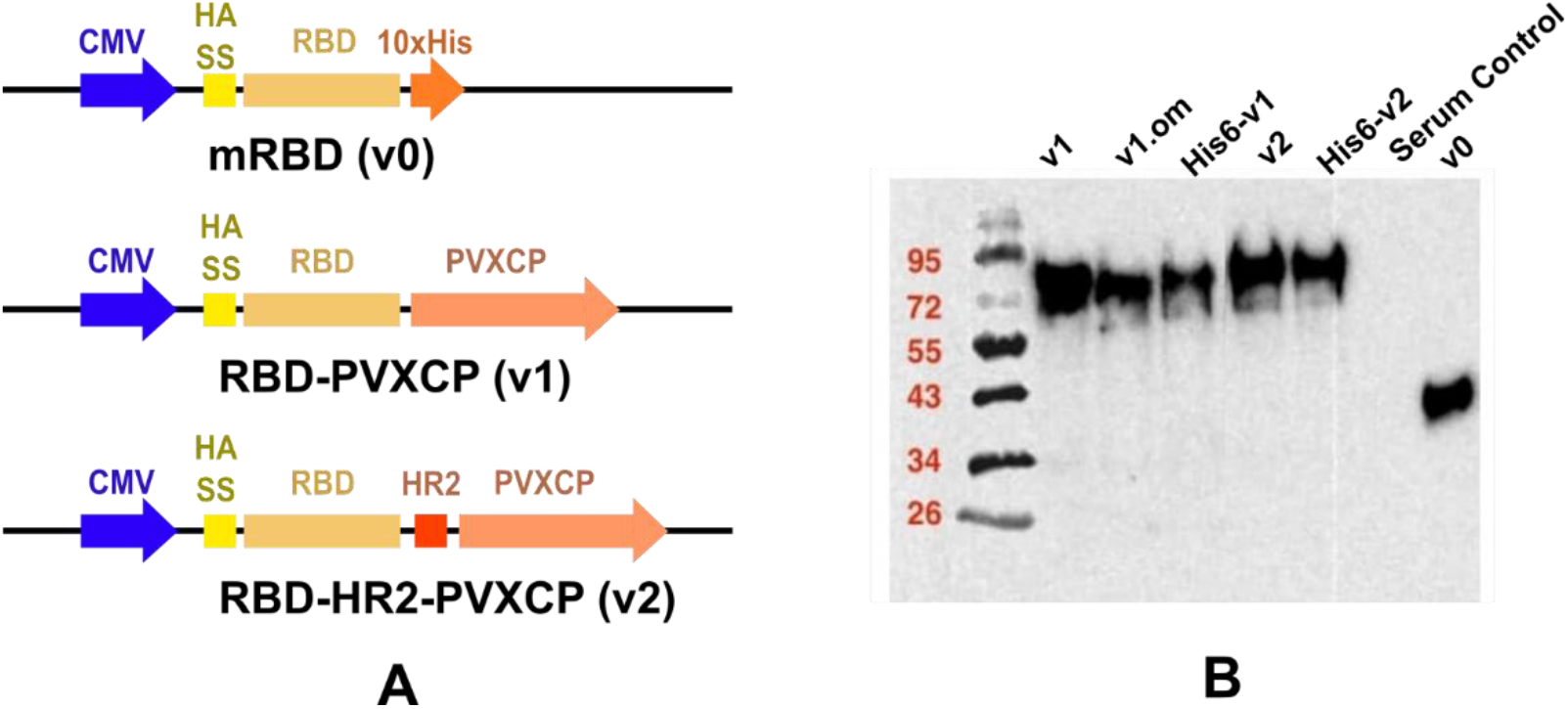
Schematic depiction of DNA vaccine constructs (A) and WB confirmation of secreted proteins in HEK293F culture supernatants (B).

In order to assess the RBD-HR2-PVXCP fusion vaccine protein and its behavior in a water solvent, 500 ns Molecular Dynamics (MD) simulation in the explicit solvent was carried out. Figure 2 illustrates the 3D structure of the RBD-HR2-PVXCP protein and its multimeric state after the oligomerization with the expected diameter of the molecule.

**Figure 2.**
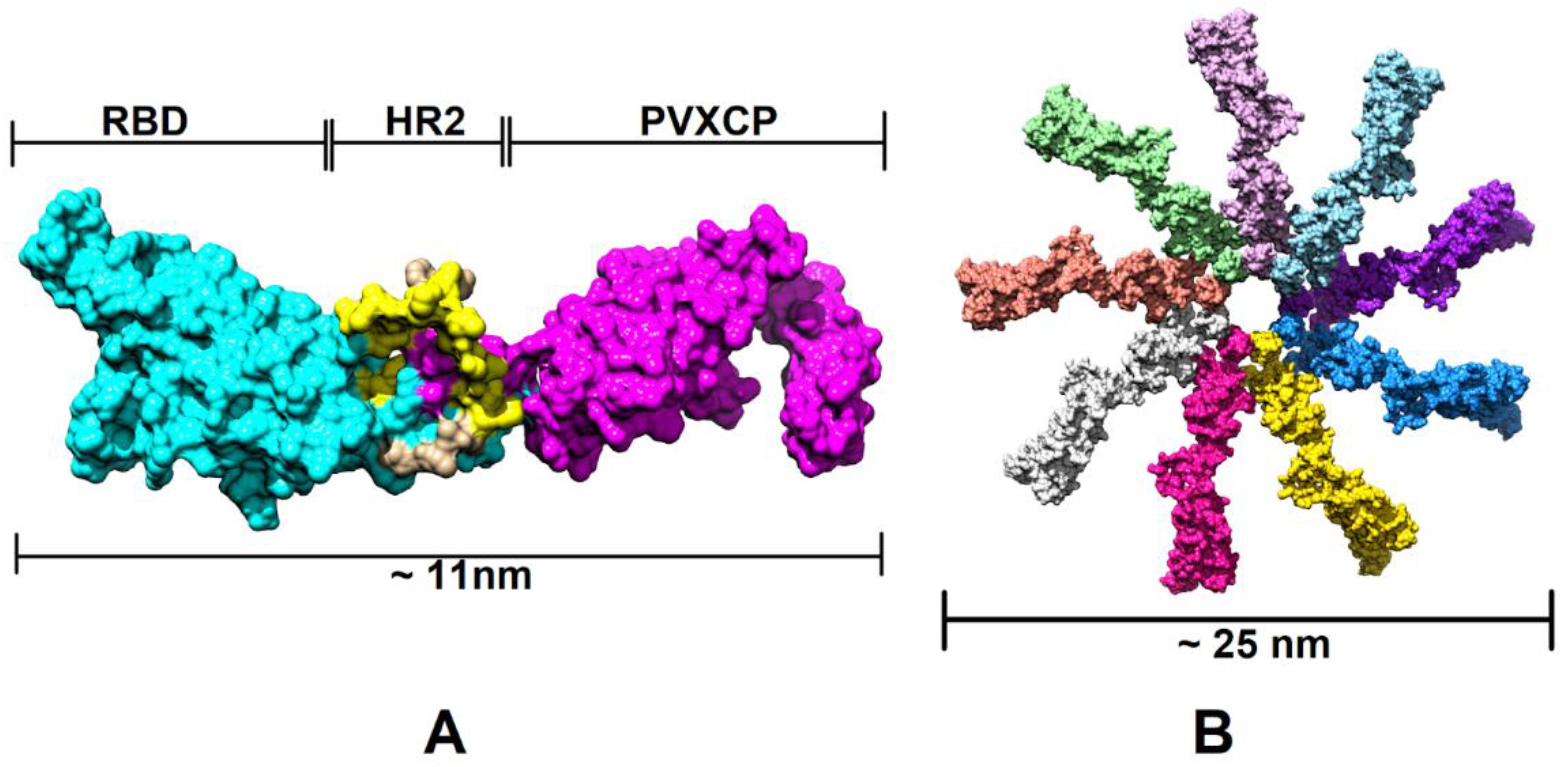
Structural illustration of the RBD-PVXCP vaccine, based on the RBD (PDB: 7KLW), and PVXCP (PDB: 6R7G) structures (A) and its 9-mer structure (B). Illustrated size of the protein complex was measured in UCSF Chimera software.

Western blot analyses with the anti-S-protein polyclonal sera confirmed soluble expression of vaccine candidates in cultural medium after the transient transfection of HEK293 FreeStyle (Figure 1B). The molecular weights of the proteins corresponded to the theoretically predicted taking into account glycosylation of RBD. The expression level of PVXCP fusion proteins was measured at 5 ug\ml level which is not inferior to the monomeric RBD transient transfection reported elsewhere^51^.

In order to determine RBD-PVXCP fusion protein oligomerization status, a hexahistidine (His-6) tag was added to the N-terminal part of constructs v1 and v2 between the RBD and HA-signal sequence. These fusion proteins were purified from 100 ml HEK293 FreeStyle supernatant by means of metal-chelating affinity chromatography (Ni-NTA) and were analyzed using dynamic light scattering (DLS). It was found that the majority of the proteins (>97% for RBD-PVXCP and >85% for RBD-HR2-PVXCP) had a radius of 3.5 – 12.7 nm (Figure S2). A weighted average radius of 11.3 nm for RBD-HR2-PVXCP corresponds to the size ranging between a PVXCP disk (Fig. 2B) or two PVXCP disks stacked together^52^.

This result allows us to suggest the multimerization effect of PVXCP protein in addition to its adjuvant properties. This also corresponds well to the generated molecular model of RBD-HR2-PVXCP monomer and oligomer.

### 3.2 Humoral response of DNA vaccine candidates in mice

The immunogenicity of the candidate DNA vaccines in mice was assessed in two experiments. First, we compared RBD-specific IgG serum levels of mice immunized with DNA vaccine v1, v2 or empty pIF vector. Three groups of female BALB/C mice (6 animals per group) were immunized twice (with the 14 days interval) with 2.5 μg of pDNA followed by electroporation at the injection site (Figure 3A). Two weeks after boost immunization blood was collected from all animals to evaluate humoral response. All animals were sacrificed to obtain spleens for cellular response analysis.

**Figure 3.**
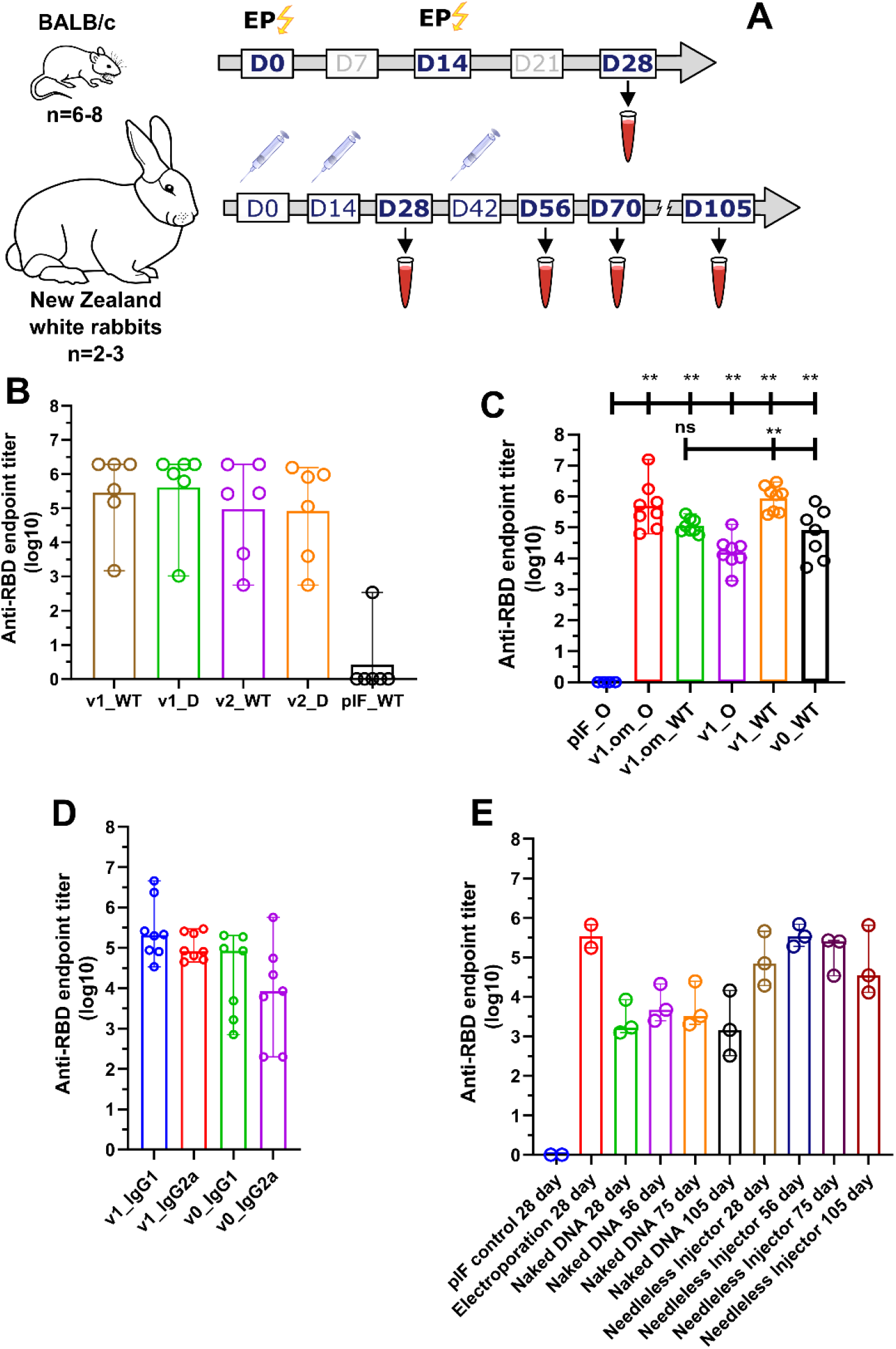
Analysis of RBD-specific humoral response. (A) Schematic representation of the experiment schedule; (B) RBD-specific IgG antibody titers from mice immunized with 2.5 ug pDNA; (C) RBD (omicron and wild-type)-specific IgG antibody titers from mice immunized with 50 ug pDNA. ns - p value > 0.05, ** - p value <0.01 (Mann Whitney U-test); (D) Anti-RBD antibodies isotype analysis; (E) RBD-specific IgG antibody titers from rabbits immunized with 100 ug pDNA by various routes.

Following immunization, no local inflammation response at the injection site or other adverse effects were observed during the observation period. A prime\boost immunization of v1 and v2 induced production of SARS-CoV-2 RBD-specific IgG antibodies with the mean endpoint titer approached 287 424 and 94 294 respectively two weeks after the last immunization (Figure 3B). We also evaluated serum IgG binding to B.1.617.2 (Delta) variant with the resulted mean endpoint titers 409 129 and 82 001 for v1 and v2, respectively.

In order to unleash the full antigenic potential of v1 - 50 μg pDNA dose was used for the immunization as well as v1.om variant for the emerging B.1.1.529 SARS-CoV-2 variant. Adjuvant properties of PVXCP were assessed by the comparison of the humoral response to v1 and v1.om vaccines with the vaccine that encoded monomeric RBD (v0) without fusion partner (Figure 3C). Calculated mean endpoint titers of collected sera against Wuhan-Hu-1 RBD (WT) and B.1.1.529 (Omiron) are presented in Table 1.

**Table 1.** RBD-specific IgG antibody mean endpoint titers from mice immunized with 50 ug pDNA

| RBD\Vaccine | v0 | v1 | v1.om |
| --- | --- | --- | --- |
| WT | 62 010 | 869 124 | 113 654 |
| Omicron | NA | 16 053 | 497 209 |

Taking into account, that ELISA antibody titres typically correlate with the neutralizing antibody titres, we have utilized ELISA to compare different design’s IgG levels. It was found that the addition of PVXCP leads to more than a tenfold increase in anti-RBD titers. Moreover, binding of serum IgG from mice immunized with the v1.om to wild-type RBD were only five times lower in comparison with the Omicron RBD binding, while binding of IgG from mice immunized with the v1 were fifty time lower for Omicron RBD in comparison with wild-type RBD. This suggests that RBD-PVXCP vaccine with the RBD harboring Omicron mutations could be referred to as a more promising anti-SARS-CoV-2 vaccine candidate.

We determined the IgG subclasses of RBD-specific antibodies induced by the v0 and v1 by sandwich ELISA (Figure 3D). Isotype analysis indicated that the binding antibodies elicited by monomeric RBD was IgG1 > IgG2a that indicates a strong Th2 dominant immune response ^53^. The addition of PVXCP in the v1 design allowed the leveling of IgG1 and IgG2a antibodies, which indicates a bias to the more preferred Th1 immune response.

### 3.3 A DNA vaccine-induced T cell response in mice

In the first experiment (vaccination with 2.5 μg of pDNA) T cell response was evaluated by ELISpot measurement of IFNγ production by splenocytes after S1 peptide pool stimulation. Both variants of v1 and v2 vaccines demonstrated significant response. A similar pattern of vaccine immunogenicity was observed for the spot number and spot size in the ELISpot test (Figure 4).

**Figure 4.**
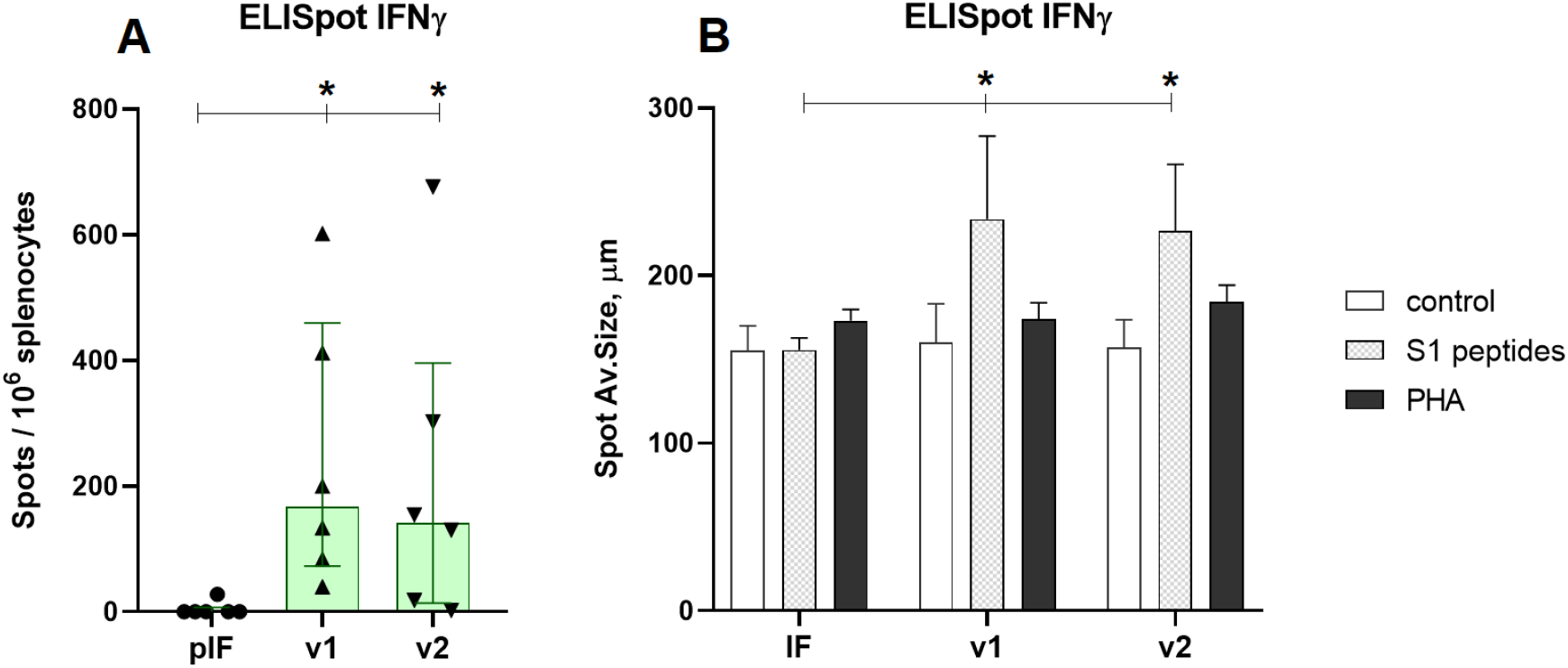
Analysis of RBD-specific cellular immune response to low-dose vaccines. A - Spots count for IFNγ, B - Spots size for IFNγ. pIF - vector control, v1 - RBD-PVXCP, v2 - RBD-HR2-PVXCP vaccines * indicates the significance of differences (p<0.05) (Mann Whitney U-test).

In the second experiment with 50 μg of pDNA vaccines T cell response was estimated by ELISpot measurement of IFNγ and IL-4 production. The aim of the experiment was to compare the maximum immunogenicity potential of the v1 vaccine and v1.om against RBD monomer (v0). The production of IFNγ in response to antigen stimulation in mice vaccinated with all three vaccine variants was significantly higher than the control vector, expressed both in spots count and average spots size (Figure 5A,C). The highest response was to v1, but the difference between vaccine variants was not significant (v0 vs v1, p=0.23).

In each experimental group there were 2-3 mice that responded to vaccination with IL-4 production, but the differences between vaccines and vector controls were not significant, and the intensity of the IL-4 response was significantly lower than IFNγ response (Figure 5B). The ratio IFNγ/IL-4 was 5,3 for v0 (p=0.011), 48.3 for v1 (p<0.001), and 8.9 for v1.om (p=0.013).

**Figure 4.**
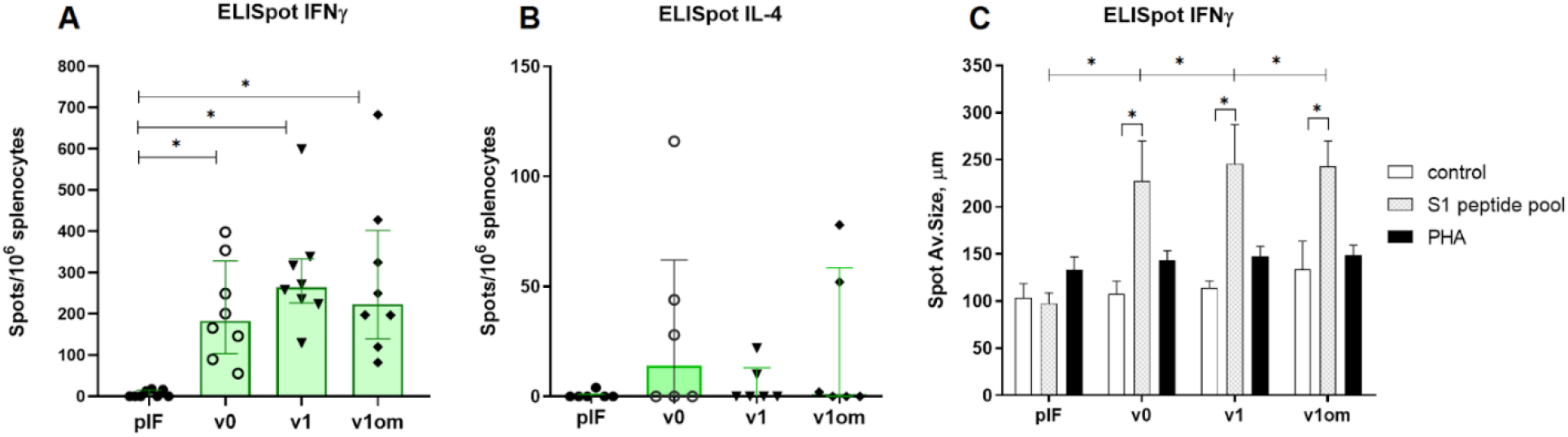
Analysis of RBD-specific cellular immune response to high-dose vaccines. A - Spots count for IFNγ, B - Spots count for IL-4, C - Spots size for IFNγ. v0 - monomer RBD (wt), v1 - RBD(wt)-PVXCP, v1om - RBD(omicron)-PVXCP * indicates the significance of differences (p<0.05) (Mann Whitney U-test).

### 3.4 A Needle-Free injection delivery of RBD-PVXCP DNA-vaccine in rabbits

We next evaluated immunogenicity of v1 anti-SARS-CoV-2 vaccine candidate in rabbits, an animal model well suited for immunological and toxicological studies^54^. As the electroporation is a traumatic and painful procedure with very few apparatus options, and naked pDNA delivery in aqueous solution lack of efficiency, we decided to explore a more robust and clinically tested approach - needle-free injection^55^.

We have compared three delivery routes for v1 pDNA delivery - EP (Group 2), needle injection of naked DNA (Group 3) and needle-free injection (Group 4). We performed prime\boost immunization in a two week interval (day 0, day 14) and the second boost for naked DNA and needle-free injection (day 56). The second booster immunization for electroporation was not performed due to the traumatic impact of this method on rabbits. The electroporation (Group 2) in our hands caused a local skin-deep burn. The jet-injection in Group 4 was well tolerated and resulted only in a hematoma in some cases at the injection site that were almost fully recovered within 2 weeks after immunization. The needle-injection (Group 1 and 3) was well tolerated and had no obvious local side-effects.

The sera from the vector control (pIF) Group 1 produced only the background level of absorbance signal. The v1 pDNA vaccine induced anti-RBD-specific immune response in all routes of administration. At day 28 (week 4), the mean RBD-specific end-point titer were 343 887, 2 614 and 85 712 for EP, naked DNA and needle-free injection respectively (Figure 3E). A second boost of naked DNA and needle-free injected DNA resulted in the mean endpoint titer increasing to 6 274 and 352 906 at the day 56. It is worth noting that after third immunizations by needleless injection, the antibody titer matched the titer after electroporation in a prime\boost regimen.

Observation of Group 3 and Group 4 of rabbits after immunization was continued and two more blood samples (at day 75 and at day 105) were collected. Endpoint titer in Group 4 decreased to 132 882 at day 75 and to 66 633 at day 105.

## 4 Discussion

The COVID-19 pandemic is the first worldwide challenge of this scale since H1N1 Spanish flu^56^. It greatly accelerated novel vaccines and therapeutic antibodies discovery, development and deployment. Novel platforms were approved for clinical uses including mRNA, DNA and vector 57 vaccines^57^.

The global vaccine campaign also provides a unique opportunity to compare different platforms and technologies at an unprecedented scale. It has become apparent that mRNA and vector vaccines have surpassed traditional inactivated and recombinant protein vaccines in a term of efficiency^58^. Despite that more than 15 DNA vaccine candidates entered different stages of clinical trials, only ZyCoV-D has been approved in a single country (https://covid19.trackvaccines.org/). Based on the results of a Phase III clinical trial, ZyCoV-D SARS-CoV-2 DNA vaccine has been found to be 67% protective against symptomatic COVID-19 delivered by needle free injection device. India’s drug regulator has approved it as a first DNA vaccine for humans for prophylaxis against COVID-19. However, its efficiency is significantly lower than for approved mRNA vaccines Comirnaty and Spikevax raising question about similar DNA-vaccines worldwide deployment^59,60^.

Here, we approached SARS-CoV-2 DNA-vaccine development with a rational design strategy of immunogen molecules. We developed two variants of SARS-CoV-2 vaccine based on the fusion of RBD immunogen with the Potato virus X coat protein (PVXCP) adjuvant molecule in order to enhance the depth and quality of the immune response to RBD.

PVXCP could self-assemble into virus-like particles even in the absence of Potato Virus X RNA, enhancing the depth of immune responses to cancer DNA vaccines^52^. Displaying antigens on nanoparticles or virus-like particles is an established strategy to increase immunogenicity. Multivalent antigens that can mimic the repetitive and well-ordered antigenic structures found on many pathogens cross-link B-cell receptors and activate B-cells more efficiently than their monovalent counterparts. In addition, they can be taken up by antigen-presenting cells and trafficked to lymph nodes more efficiently, leading to improved formation of germinal centers^61,62^.

Different nanoparticle vaccines against SARS CoV-2 have been developed and revealed an advantage over monovalent counterparts^63,64,65,66^. PVXCP could tolerate long polypeptides fused to its N-terminus, maintaining high protein expression. PVXCP is also described as an immune enhancer that stimulate CD4 help and shift immune response towards Th1^67^.

Our data support the high level of the secretion of v1 and v2 vaccine candidates in HEK293F system. It was revealed that PVXCP fused to the RBD with or without the HR2 region between them does not affect RBD expression level. Purified RBD-PVXCP and RBD-HR2-PVXCP proteins encoded by v1 and v2 also tends to oligomerize in solution. This, if applicable under *in vivo* conditions should result in stronger humoral response in comparison with the monomeric RBD protein.

In mice, strong humoral and T cell responses were observed using 2.5 ug and 50 ug pDNA v1 and v2 electroporation. Anti-RBD IgG titers exceeded 10^5^ even for smaller 2.5 ug dose and what is more important - serum IgG binding to RBD was not affected by the presence of B.1.617.2 mutations in RBD. We also evaluated humoral response elicited by v1.om variant and compared it to the v1 and v0. The addition of PVXCP to RBD elevated RBD-specific IgG more than tenfold for the wild-type RBD. As a result we observed the immunogenicity not inferior and sometimes superior to the preclinical data of other DNA-based and mRNA-based vaccines^68,69,70^.

Surprisingly, omicron-based v1.om vaccine outperformed wild-type-based v1 vaccine by eliciting more universal antibodies, which are more tolerant to RBD-mutations.

We found a prominent T-cell immune response expressed in IFNγ production against the S1 peptide library in vaccinated mice. The highest response rates of 700 with median 200-300 spots per 10^6^ splenocytes were observed for both 2.5 μg and 50 μg doses of v1 and v2 vaccine candidates. This response rate exceeds the results of even the higher dose of 100 μg DNA vaccine ZyCoV-D and approaches the response rate of vaccine INO-4800 and pGO-1001/1002, which encodes the whole Spike protein as an antigen ^49, 71, 72^. In the same experiment, an immune response to the vaccine expressed in IL-4 production was observed in 33-50% of mice at a level several times lower than IFNγ production.

Immunization of children with whole-inactivated-virus vaccines against RSV and measles virus led to vaccine-associated enhanced respiratory disease (VAERD), an immunopathological state associated with T helper 2 cell (TH2)-biased immune responses^73,74^. A similar phenomenon has been seen in some animal models with whole-inactivated vaccines, including SARS-CoV vaccines^75,76^. The significance of the Th1 cell response has also been shown for asymptomatic and mild forms of COVID-19 infections^77,78^.

A higher IFN/IL-4 ratio in our experiments indicates the development of a Th1 type immune response in BALB /c mice naturally biased for Th2 immune response. This is presumably due to the role of the PVXCP protein, since the vaccine with the RBD monomer induces a more apparent production of IL-4.

We also observed PVXCP-induced Th1 help forwarded bias by comparing the isotypes of antibodies induced by DNA-encoded RBD vaccine v0 and RBD-PVXCP v1. It was revealed that IgG1>>IgG2a antibodies levels for v0 shifted to IgG1~Ig2a ratio for v1. It confirms our assumption that PVXPC could turn anti-RBD immune response to a more favorable Th1 type.

RBD-PVXCP vaccine also induced a strong humoral response in rabbits after prime\boost immunization. Since electroporation has limited use in humans, especially for prophylactic vaccination we focused on needle-free injection systems, that a clinically approved, cheap and deliver the payload with high efficiency without considerable side effects.

After the prime/boost set of our pDNAv1 vaccine the mean RBD-specific end-point titer reached 85 712 utilizing needle-free injection method. To the best of our knowledge – we obtained the highest IgG level among all the naked pDNA vaccines that were pre-clinically tested in rabbits without non-applicable on humans electroporation ^40, 49^. In order to further explore the potential of our vaccine candidate at week 6 we performed a second boost that raised the mean endpoint titer to 352 906 approaching electroporation in the prime\boost regimen. Two months after the second boost IgG level still was high, with the end-point titer 66 633. It confirms the long-lasting immunity induced by RBD-PVXCP vaccine candidate.

Summarizing all of the above, DNA vaccine candidates that were rationally designed and developed in the present work, demonstrate high immunogenicity and good safety profile in two animal models - mice and rabbits. Evaluation of a neutralization potency of elicited antibodies and the prophylactic effect in SARS-CoV-2-challenged animal models is needed.

In conclusion, our current work demonstrates the feasibility and potency of SARS-CoV-2 DNA vaccines to induce anti-RBD IgG titer compared with mRNA-LNP vaccines, providing important information for further development of DNA-based vaccines even beyond SARS-CoV-2 infection.

## Acknowledgments

The authors express their gratitude to the Institute of Physiology of the National Academy of Sciences of Belarus for the assistance in keeping laboratory animals.

## Author Contributions

Conceptualization and design of the experiments: DD, AMe and MK. Vaccine candidates design and expression analysis: DD, MSh, SD and AK. MD simulation and statistical analysis: MSh. Vaccine candidates production - SG and DL. Animal experiments and ELISpot analysis: AMe, MSt, SG. Drafting the manuscript: DD, AMe and MK. All authors contributed to the article and approved the submitted version.

## Conflict of Interest

DD, AMe and MK have a pending patent application for the DNA-based SARS-CoV-2 vaccine. The other authors declare no competing interests.

## Funding

The authors declare that this study received funding from Immunofusion. The funder was not involved in the study design, collection, analysis, interpretation of data, the writing of this article or the decision to submit it for publication.

