## Supplementary material for "Design and Immunogenicity of SARS-CoV-2 DNA vaccine encoding RBD-PVXCP fusion protein": Figure S1, Figure S2

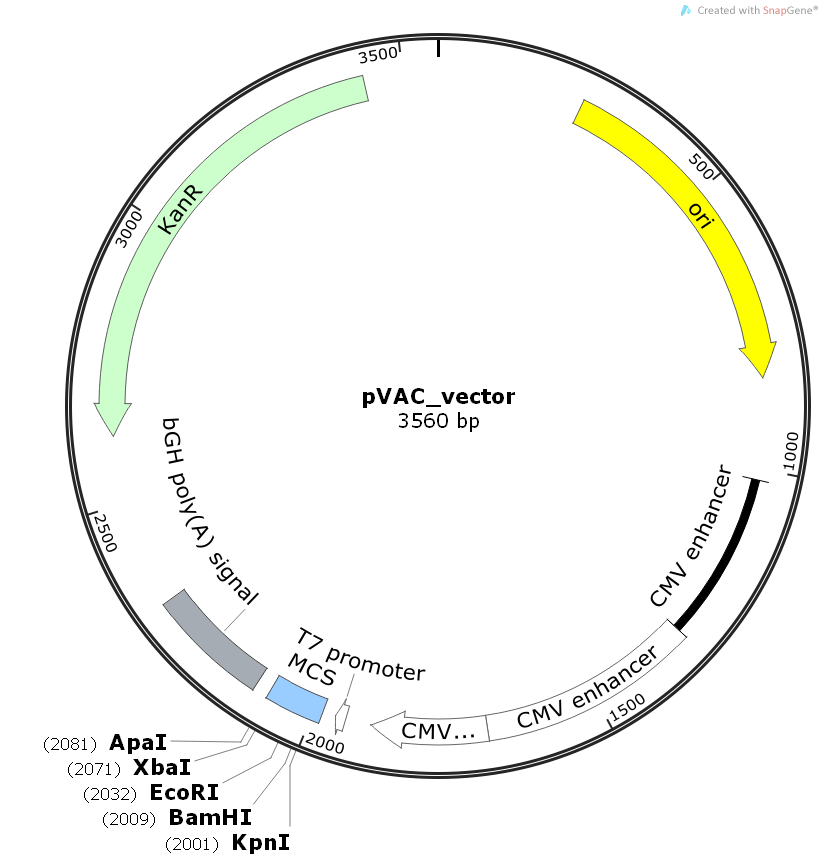
**Figure S1. Generic plasmid vector map for the pIF DNA-vaccine delivery vector**

**Figure S2. Comparing histograms of the DLS-measured particles distribution.**

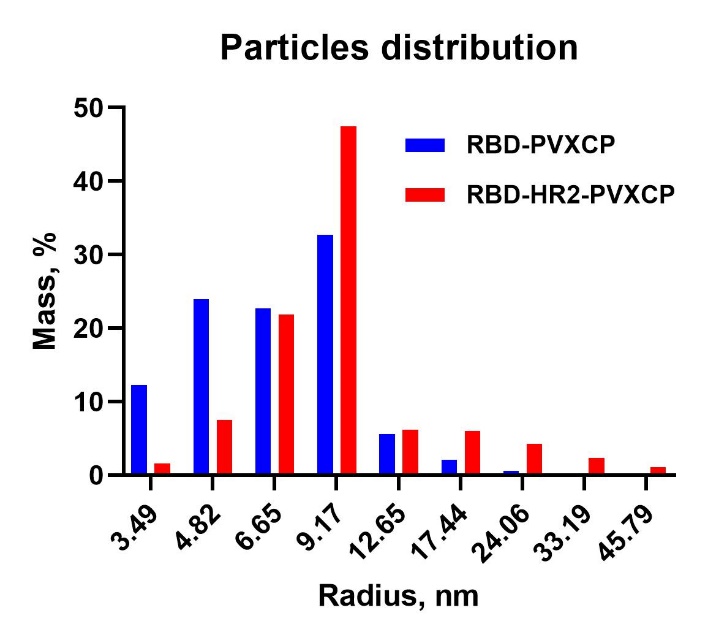
